# Research gaps and new insights in the intriguing evolution of *Drosophila* seminal proteins

**DOI:** 10.1101/2021.05.11.443674

**Authors:** J Hurtado, FC Almeida, SA Belliard, S Revale, E Hasson

**Affiliations:** Departamento de Ecología, Genética y Evolución, Facultad de Ciencias Exactas y Naturales, Universidad de Buenos Aires (UBA), CABA, Argentina; Instituto de Ecología, Genética y Evolución de Buenos Aires, Consejo Nacional de Investigaciones Científicas y Técnicas (CONICET), CABA, Argentina; Laboratorio de Insectos de Importancia Agronómica, IGEAF (INTA), GV-IABIMO (CONICET), Hurlingham, Buenos Aires, Argentina; Wellcome Trust Centre for Human Genetics, University of Oxford, Oxford, OX3 7BN, UK

**Author notes:** Author for Correspondence: Juan Hurtado.

**Keywords:** Accessory glands, *De novo* gene evolution, Gene co-option, Gene turnover, Gene origin, Seminal fluid

## Abstract

While the striking effects that seminal fluid proteins (SFPs) exert on females are fairly conserved among Diptera, they exhibit remarkable evolutionary lability. Consequently, most SFPs lack detectable homologs among the repertoire of SFPs of phylogenetically distant species. How such a rapidly changing proteome “manages” to conserve functions across taxa is a fascinating question. However, this and other pivotal aspects of SFPs’ evolution remain elusive because discoveries on these proteins have been mainly restricted to the model *D. melanogaster*. Here, we provide an overview of the current knowledge on the inter-specific divergence of *Drosophila* SFPs and compile the increasing amount of relevant genomic information from multiple species. Capitalizing the accumulated knowledge in *D. melanogaster*, we present novel sets of high-confidence SFP candidates and transcription factors presumptively involved in regulating the expression of SFPs. We also address open questions by performing comparative genomic analyses that failed to support the existence of conserved SFPs shared by most dipterans and indicated that gene co-option is the most frequent mechanism accounting for the origin of *Drosophila* SFP-coding genes. We hope our update establishes a starting point to integrate, as more species are assayed for SFPs, further data and thus, to widen the understanding of the intricate evolution of these proteins.

## Introduction

During mating, spermatozoa expelled from the testes travel throughout the ejaculatory duct into the female reproductive tract accompanied by a rich repertoire of proteins and peptides known as Seminal Fluid Proteins (SFPs) (reviewed in, e.g., Avila et al. 2011; Avila et al. 2016; Chapman 2008; Ramm 2020). These proteins, likely adapted to sperm competition and fertilization, have been highly studied in *Drosophila melanogaster* (e.g., Civetta & Ranz 2019; Hopkins, Sepil, Bonham et al. 2019; Hopkins, Sepil, Thézénas et al. 2019; Misra & Wolfner 2020; Ravi Ram et al. 2005; Ravi Ram & Ramesh 2003; Ravi Ram & Wolfner 2007; Wigby et al. 2020; Wolfner 2007). Once inside the female, some of these proteins will remain bound to spermatozoa, contributing to sperm functions, and some may even interact with the already stored sperm from previous mates (e.g., Avila et al. 2011; Holman 2009; Misra & Wolfner 2020; Ravi Ram & Wolfner 2007; Singh et al. 2018; Wolfner 2007). Many others instead will interact intimately with female biomolecules in the reproductive tract and other organs, and are capable of changing drastically her physiology and behavior (e.g., Avila et al. 2011; Avila et al. 2016; Avila & Wolfner 2017; Lung & Wolfner 1999; Ravi Ram et al. 2005; Ravi Ram & Wolfner 2007).

In *Drosophila*, decrease of female receptivity to mating, increase of egg production, and conformational modification of the female reproductive organs stand out among the profound changes that SFPs trigger in the female (reviewed in Avila et al. 2016). Given the conflicts of interest between males and females (and between competing males), some of the SFPs effects, while beneficial to the last-mating male, can be detrimental to the female (Chapman et al. 1995; Lung et al. 2002; Mueller et al. 2007; Wigby & Chapman 2005). Thus, rapid antagonistic coevolution is expected between some SFPs and female-derived proteins that interact with them (e.g., Sirot et al. 2014). Nevertheless, other SFPs work synergistically with female biomolecules to facilitate fertilization or progeny production for the mutual benefit of males and females (Avila et al. 2016; Wolfner 2009). Therefore, they are expected to diverge more slowly. In fact, sequence comparisons between closely related *Drosophila* species revealed that some SFPs have evolved extremely fast by positive selection while others are conserved by purifying selection (e.g., Almeida & Desalle 2008; Haerty et al. 2007; Turner & Hoekstra 2008; Wong et al. 2012).

The biochemical classes into which SFPs typically fall (e.g., proteases, protease inhibitors, lectins, lipases, and cysteine-rich secretory proteins) seem quite conserved among Diptera, even among animals from different classes (reviewed in, e.g., Avila et al. 2016; Wigby et al. 2020). This suggests that the functional spectrum of SFPs is adaptively restricted at the molecular level. Nonetheless, a striking pattern for the vast majority of SFPs is the lack of detectable homologs among SFPs of phylogenetically distant species (Ahmed-Braimah et al. 2017; Almeida & Desalle 2009; Davies & Chapman 2006; Haerty et al. 2007; Mueller et al. 2005). Even though the rapid divergence of some of these proteins may hinder homology detection, the main reason behind this pattern seems to be the rapid turnover (gain and loss) of genes encoding SFPs (seminal genes) (Sirot 2019; Sirot et al. 2014). It remains unknown, however, whether a core of a particular SFPs, playing essential reproductive roles, has been conserved over long evolutionary periods. Neither do we know how new seminal genes arise so frequently, or to what extent regulatory elements of seminal genes are conserved across species.

Addressing these broad evolutionary questions requires performing multi-species comparative analyses which, in turn, requires extensive omic information on the seminal proteome of several related species. While most of the achieved findings on SFPs have been restricted to *D. melanogaster*, in recent years, the seminal proteome has been characterized in many other species, including Drosophilids. This brings up an opportunity to use the *Drosophila* model to address open questions on SFPs evolution and capitalize the accumulated knowledge in *D. melanogaster*.

Here, to elucidate some answers, we review the current knowledge on the evolution of *Drosophila* SFPs, compiled genomic data from multiple species, and performed molecular evolutionary analyses using bioinformatic tools. We structured the text into sections, each of which tackles a specific topic by presenting knowledge gaps, new insights, and future perspectives.

## Identification

In *D. melanogaster*, as in many other dipteran species, the main secretory tissues of the male reproductive system are the accessory glands, a pair of merocrine glands attached to the anterior region of the ejaculatory duct (Avila et al. 2016; Chen 1984; Gillott 1996). While mutant males without accessory glands cannot elicit the normal postmating responses in their female mates (Kalb et al. 1993), it has long been known that ACcessory glands Proteins (ACPs) alone are sufficient for triggering these responses in virgin females (reviewed in Ravi Ram & Wolfner 2007). In fact, the first studies on male reproductive proteins aimed to identify SFPs focusing on the male accessory glands.

The very first SFP to be identified was ‘Sex Peptide’ (SP, also known as Acp70A). It was purified from an HPLC fraction of accessory gland extracts that proved, after being injected into virgin females, to reproduce the well-known postmating responses (Chen et al. 1988). The authors also showed that SP gene is transcribed specifically in the male accessory glands. Afterwards, diverse methods such as Expressed Sequence Tags screening, RT-PCR, subtracting hybridization, and cDNA microarray hybridization allowed the identification of many other genes specifically expressed in the male accessory glands (reviewed in Chapman & Davies 2004). Among those genes, the ones encoding proteins or peptides with a predicted signal peptide—that permits canonical merocrine secretion—were considered as candidate seminal genes (Ravi Ram & Wolfner 2007; Swanson et al. 2001). By 2005, using this double criterion, accessory gland-specific expression and capacity to encode secretory proteins, it was possible to identify ~90 putative seminal genes. Five additional seminal genes—or presumptive seminal genes—were found in other organs of the male reproductive tract: the testes, the ejaculatory duct, and the ejaculatory bulb (Cavener & MacIntyre 1983; Dyanov & Dzitoeva 1995; Kopantseva et al. 1990; Ludwig et al. 1991; Lung & Wolfner 2001; Richmond et al. 1980; Saudan et al. 2002; Sheehan et al. 1979). Seven additional candidate genes were identified by mass spectrometry of tryptic peptides from accessory glands secretions (Walker et al. 2006).

Until 2008, only 22 of the predicted seminal genes were confirmed, mainly by means of immunological techniques, to be transferred to females during mating (e.g., Aigaki et al. 1991; Bertram et al. 1996; Cho et al. 1999; Coleman et al. 1995; Kopantseva et al. 1990; Lung & Wolfner 1999, 2001; Meikle et al. 1990; Ravi Ram et al. 2005; Wong et al. 2008). In 2008, Findlay et al. conducted a proteomic screen that largely extended the list of proven SFPs. The authors used isotopic labeling of the female to distinguish, among proteins isolated from the reproductive tract of newly mated females, between female proteins and proteins transferred from unlabeled males. In this way, they confirmed 75 of the previously predicted SFPs and revealed 63 novel ones. More recently, Sepil et al. (2019) applied quantitative proteomics to identify proteins that after mating become significantly less abundant in male reproductive tissues but more abundant in the female reproductive tract, as expected precisely for SFPs. They also cross-referenced their quantification results with transcriptomic and sequence databases to obtain a list of high-confidence candidate SFPs meeting stringent multiple criteria. Some of these candidates were already known as predicted or confirmed SFPs, while nine were novel discoveries (Sepil et al. 2019). While we were concluding this report, Wigby et al. (2020) combined data from these and other proteomic studies to provide a list of 292 *D. melanogaster* SFPs. However, the conditions they evaluated may have been too lax; according to modENCODE [implemented in FlyBase r2020_03 (Graveley et al. 2010; Thurmond et al. 2019)] and FlyAtlas2 (Leader et al. 2018), some of the genes they proposed as novel candidates are not expressed in the male reproductive tissues but in the female (e.g., *FBgn0262536, FBgn0262484*, and *FBgn0261989*), and thus, it is not clear that all these genes encode SFPs.

According to our bibliographic search, the current number of confirmed—or high-confidence candidate—non-sperm SFPs in *D. melanogaster* [hereafter Known Seminal Proteins (KSPs)] is 173 (see source studies in supplementary table S1). Our list includes 1) genes encoding proteins previously confirmed to be transferred by males to females during mating, 2) genes meeting the stringent multiple criteria adopted by Sepil et al. (2019), or 3) those genes more expressed in male reproductive tissues than in any other tissue (according to modENCODE and FlyAtlas2) also encoding secretory proteins found in the mating plug [according to Avila et al. (2015) and Wigby et al. (2020)]. Nonetheless, due to current methodological limitations, some other SFPs probably remain to be discovered. Given the leading role of accessory glands as suppliers of SFPs through merocrine secretion, genes that 1) are strongly expressed in the accessory glands and 2) encode secretory proteins can be considered seminal genes. Based on this expression/secretion (double) criterion, a suitable way of finding new candidate seminal genes may be to search in transcriptomic databases for genes expressed in the male accessory glands and to assess which of those genes encode secretory proteins using *in silico* prediction approaches.

Before searching for new candidate seminal genes, we explored to what extent *D. melanogaster* Known Seminal Genes (KSGs) meet the expression/secretion criterion by evaluating two conditions. First, we used the RNA-seq databases modENCODE (implemented in FlyBase r2020_03) and FlyAtlas2 to check which seminal genes are strongly expressed in the accessory glands. Second, we used SignalP-5.0—a deep neural network-based tool that identifies signal peptides and their cleavage sites (Almagro Armenteros et al. 2019; Nielsen et al. 1997)—to evaluate which SFPs have signal peptide required for secretion. Among the 173 KSGs, 159 (93.0%) showed relatively high expression in the accessory glands [> 25 Reads/Fragment Per Kilobase of transcript per Million mapped reads (R/FPKM), which is within the 60-70th percentile] according to one or both databases; 156 (90.2%) encoded a protein with a predicted signal peptide; 151 (87.2%) meet both conditions (supplementary table S1), and; 165 (95.3%) meet at least one of them. Most of the few genes not meeting any of these conditions are expressed specifically in the testes. These numbers not only confirm that the vast majority of SFPs are expressed in the accessory glands but also show that their secretion is mainly merocrine (but see Corrigan et al. 2014; Leiblich et al. 2012).

However, the two conditions we evaluated in the KSPs may be too lax for finding new candidate genes. For instance, accessory glands expression level could be inflated in modENCODE or FlyAtlas2, or SignalP could wrongly predict the presence of a signal peptide. Moreover, a signal peptide would only guarantee translocation into the endoplasmic reticulum followed by signal sequence cleavage. Thus, even if a gene truly meets both conditions, the protein may be retained, for instance, in the endoplasmic reticulum or the Golgi apparatus of accessory glands cells. For these reasons, we decided to evaluate *D. melanogaster* genes for a more restrictive set of six conditions that also relies on the expression/secretion criterion:

1. At least ‘Very High’ expression in the accessory glands (> 100 RPKM, which is within the ~90th percentile) according to modENCODE.
2. At least ‘Moderately High’ expression (> 25 RPKM) and expression enrichment in the accessory glands (relative to other adult tissues) according to modENCODE.
3. At least ‘Very High’ expression in the accessory glands (> 100 FPKM, which is within the ~90th percentile) according to FlyAtlas2.
4. At least ‘Moderately High’ expression (> 25 FPKM) and expression enrichment in the accessory glands (relative to whole adult male flies) according to FlyAtlas2.
5. Ability to encode a protein with a signal peptide according to SignalP.
6. Ability to encode a secretory protein according to DeepLoc, a prediction algorithm that uses deep neural networks to predict protein localization relying on sequence information (Almagro Armenteros et al. 2017). Unlike SignalP, this software differentiates between 10 subcellular localizations and distinguishes proteins of the extracellular space from proteins of the secretory pathway that are retained in the cell.

Genes fulfilling conditions 1 (or 2) and 3 (or 4) are highly (or differentially) expressed in the accessory glands according to different databases, while genes fulfilling conditions 5 and 6 are predicted to encode secretory proteins by different software programs. Therefore, we recognized 219 *D. melanogaster* genes that met conditions 1 (or 2), 3 (or 4), 5, and 6 as seminal gene candidates. These 219 genes included 122 KSGs, 43 previously predicted but unconfirmed seminal genes, and 54 newly identified candidates (fig. 1, supplementary table S1). From the 97 candidates that are not among the KSGs, 46 (22 previously predicted seminal genes and 24 novel discoveries) met all six conditions and were dubbed Unconfirmed High Confident Candidates (UHCCs) (fig. 1, table 1).

**Fig. 1.**
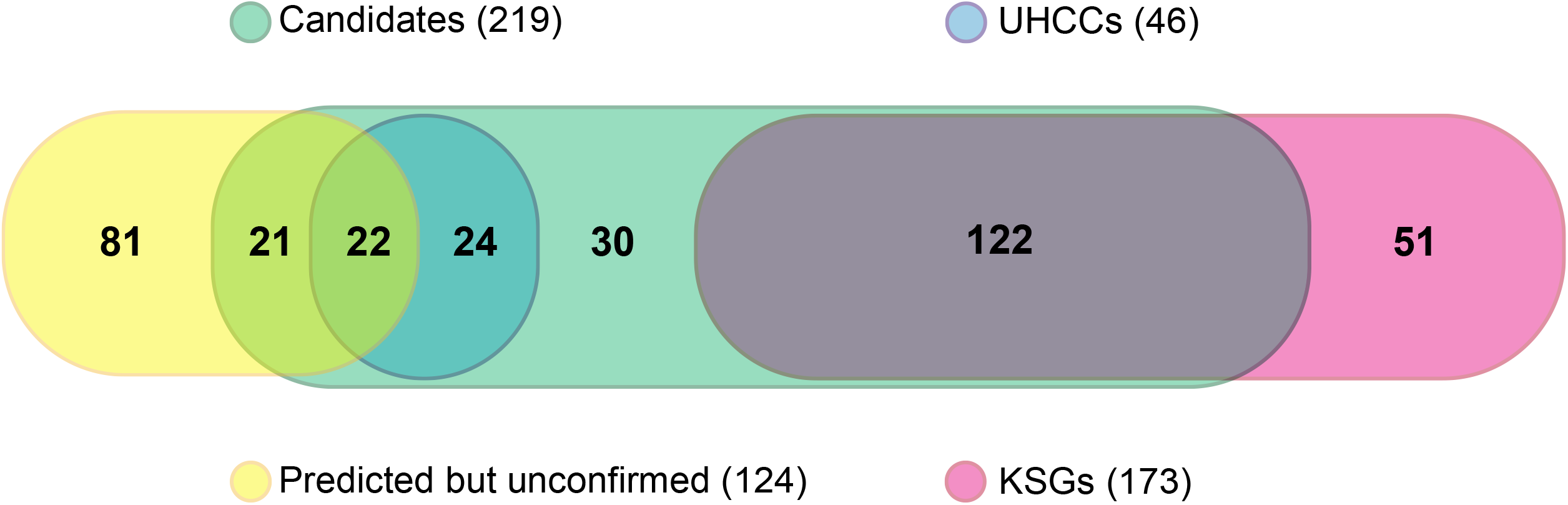
Venn diagram representing the overlap between the candidate seminal genes we identified (Candidates) and other sets of putative or confirmed *D. melanogaster* seminal genes. Candidates are those genes we identified (1) to be highly (or differentially) expressed in the accessory glands according to two transcriptomic databases and also (2) to encode secretory proteins with two software programs. Known Seminal Genes (KSGs) are those encoding proteins previously confirmed to be transferred by males into females during mating or those meeting stringent multiple criteria that indicate so. Unconfirmed High Confident Candidates (UHCCs) are those Candidates, not included among KSGs, that are both highly and differentially expressed in the accessory glands according to the two consulted transcriptomic databases. Predicted but unconfirmed seminal genes are previously predicted seminal genes not included among KSGs.

**Table 1.**
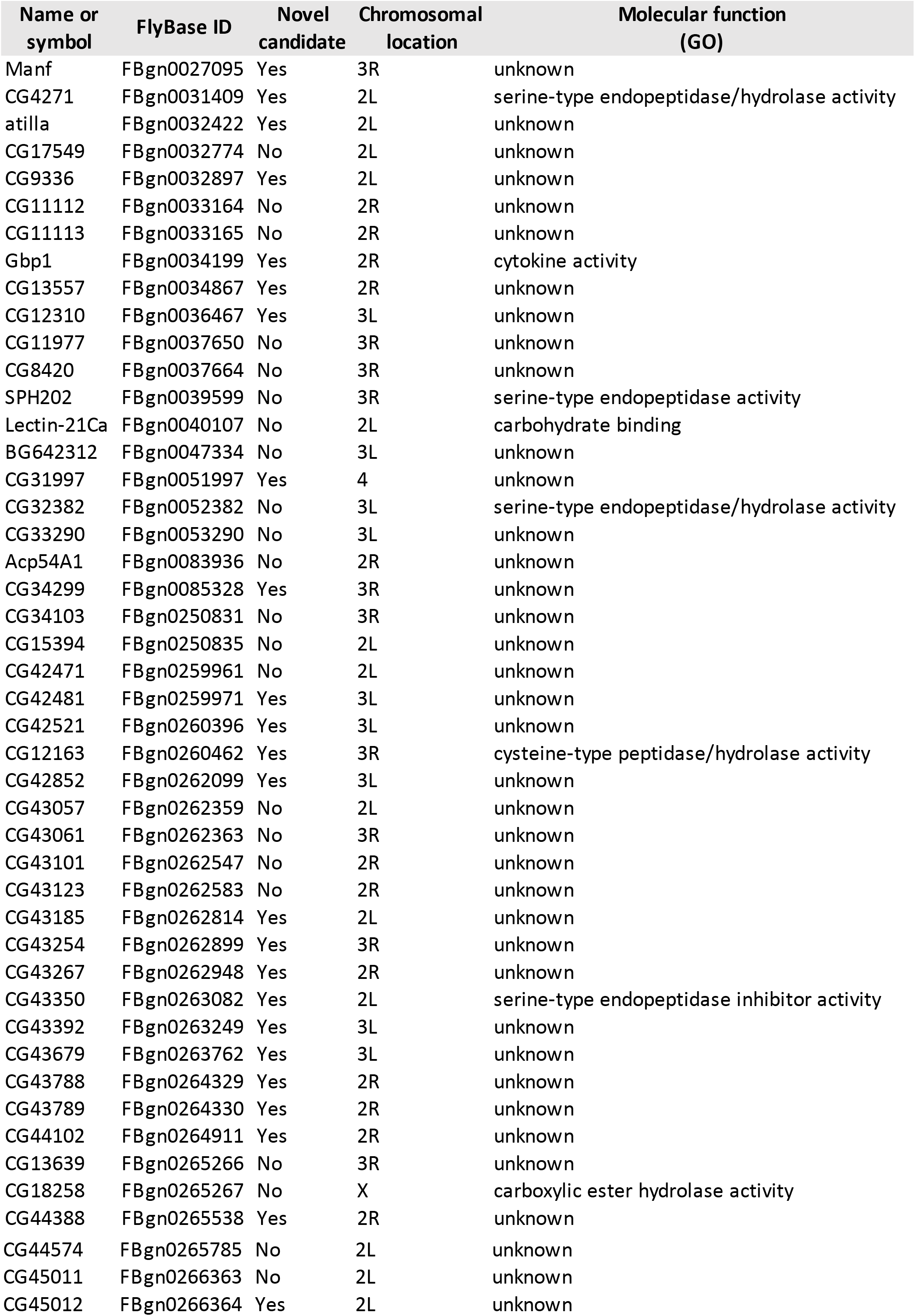
List of Unconfirmed High Confident Candidates (UHCCs). Name, chromosomal location, and molecular function (taken from FlyBase r2020_03) are shown for each gene.

| Name or symbol | FlyBase ID | Novel candidate | Chromosomal location | Molecular function (GO) |
| --- | --- | --- | --- | --- |
| Manf | FBgn0027095 | Yes | 3R | unknown |
| CG4271 | FBgn0031409 | Yes | 2L | serine-type endopeptidase/hydrolase activity |
| atilla | FBgn0032422 | Yes | 2L | unknown |
| CG17549 | FBgn0032774 | No | 2L | unknown |
| CG9336 | FBgn0032897 | Yes | 2L | unknown |
| CG11112 | FBgn0033164 | No | 2R | unknown |
| CG11113 | FBgn0033165 | No | 2R | unknown |
| Gbp1 | FBgn0034199 | Yes | 2R | cytokine activity |
| CG13557 | FBgn0034867 | Yes | 2R | unknown |
| CG12310 | FBgn0036467 | Yes | 3L | unknown |
| CG11977 | FBgn0037650 | No | 3R | unknown |
| CG8420 | FBgn0037664 | No | 3R | unknown |
| SPH202 | FBgn0039599 | No | 3R | serine-type endopeptidase activity |
| Lectin-21Ca | FBgn0040107 | No | 2L | carbohydrate binding |
| BG642312 | FBgn0047334 | No | 3L | unknown |
| CG31997 | FBgn0051997 | Yes | 4 | unknown |
| CG32382 | FBgn0052382 | No | 3L | serine-type endopeptidase/hydrolase activity |
| CG33290 | FBgn0053290 | No | 3L | unknown |
| Acp54A1 | FBgn0083936 | No | 2R | unknown |
| CG34299 | FBgn0085328 | Yes | 3R | unknown |
| CG34103 | FBgn0250831 | No | 3R | unknown |
| CG15394 | FBgn0250835 | No | 2L | unknown |
| CG42471 | FBgn0259961 | No | 2L | unknown |
| CG42481 | FBgn0259971 | Yes | 3L | unknown |
| CG42521 | FBgn0260396 | Yes | 3L | unknown |
| CG12163 | FBgn0260462 | Yes | 3R | cysteine-type peptidase/hydrolase activity |
| CG42852 | FBgn0262099 | Yes | 3L | unknown |
| CG43057 | FBgn0262359 | No | 2L | unknown |
| CG43061 | FBgn0262363 | No | 3R | unknown |
| CG43101 | FBgn0262547 | No | 2R | unknown |
| CG43123 | FBgn0262583 | No | 2R | unknown |
| CG43185 | FBgn0262814 | Yes | 2L | unknown |
| CG43254 | FBgn0262899 | Yes | 3R | unknown |
| CG43267 | FBgn0262948 | Yes | 2R | unknown |
| CG43350 | FBgn0263082 | Yes | 2L | serine-type endopeptidase inhibitor activity |
| CG43392 | FBgn0263249 | Yes | 3L | unknown |
| CG43679 | FBgn0263762 | Yes | 3L | unknown |
| CG43788 | FBgn0264329 | Yes | 2R | unknown |
| CG43789 | FBgn0264330 | Yes | 2R | unknown |
| CG44102 | FBgn0264911 | Yes | 2R | unknown |
| CG13639 | FBgn0265266 | No | 3R | unknown |
| CG18258 | FBgn0265267 | No | X | carboxylic ester hydrolase activity |
| CG44388 | FBgn0265538 | Yes | 2R | unknown |
| CG44574 | FBgn0265785 | No | 2L | unknown |
| CG45011 | FBgn0266363 | No | 2L | unknown |
| CG45012 | FBgn0266364 | Yes | 2L | unknown |

As previously noticed, *D. melanogaster* seminal genes share other quite singular features: a significantly biased location on autosomes, particularly on the second chromosome (Findlay et al. 2008; Ravi Ram & Wolfner 2007), and, on average, high *Ka/Ks* ratios (Ahmed-Braimah et al. 2017; Almeida & Desalle 2008; Haerty et al. 2007; Holloway & Begun 2004). The UHCCs resemble KSGs regarding chromosomal location (fig. 2) and *Ka/Ks* ratio (fig. 3). In addition, using the functional annotation tool DAVID (Huang et al. 2009), we performed gene-enrichment analyses for molecular function of both UHCCs and KSGs. These analyses also revealed similarities between these groups of genes: eight out of the nine (89%) Gene Ontology (GO) terms annotated to UHCCs are among the terms annotated to KSGs, and the two most represented GO terms in the UHCCs are among the over-represented terms in the KSGs (table 2). Thus, we will henceforth refer to the 173 KSGs and the 46 UHCCs together (a total of 219 genes) as an updated list of *D. melanogaster* seminal genes.

**Fig. 2.**
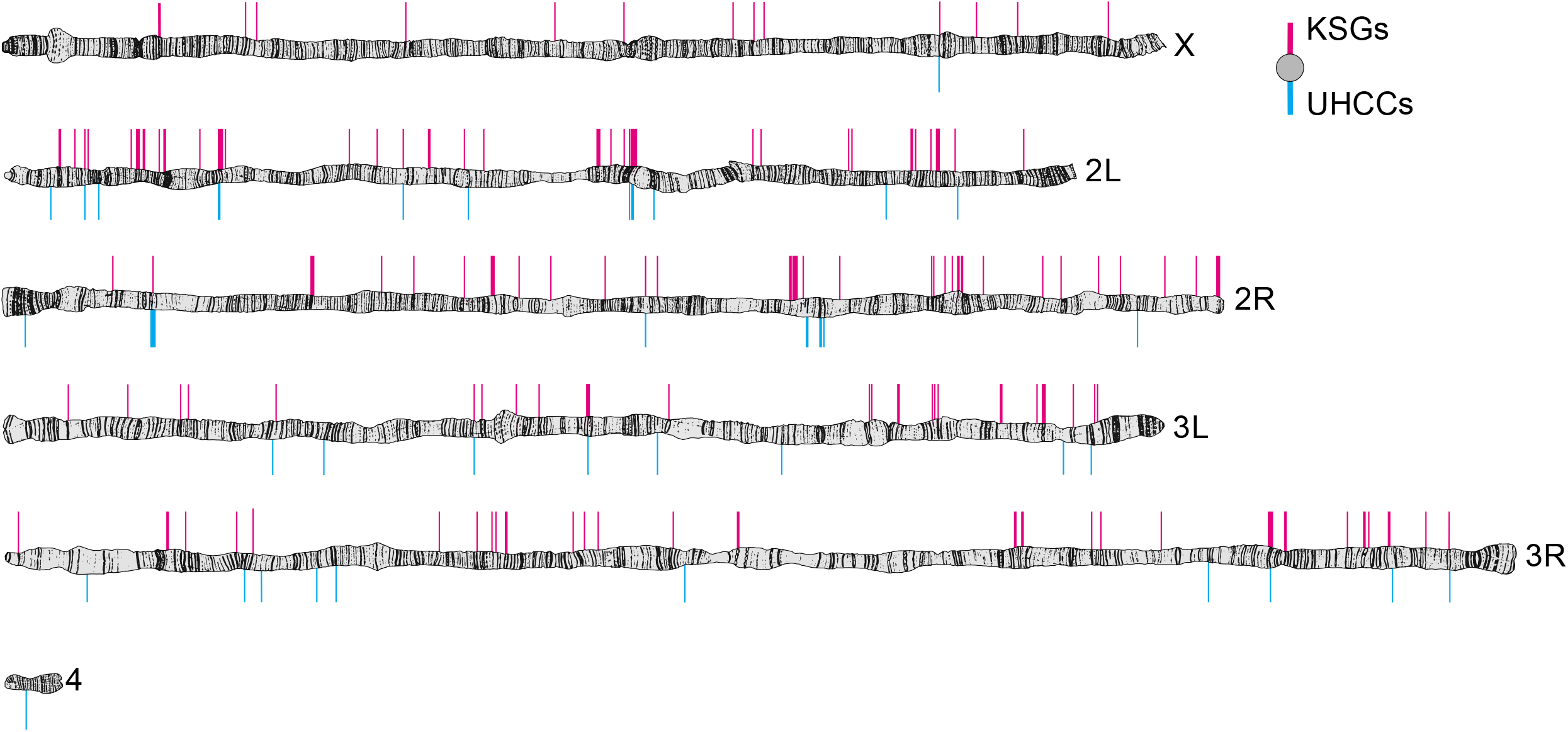
Chromosomal location of *D. melanogaster* seminal genes. Drawings of polytene chromosomes were modified from Lefevre’s photographic maps (Lefevre 1976) and gene locations were obtained from FlyBase.

**Fig. 3.**
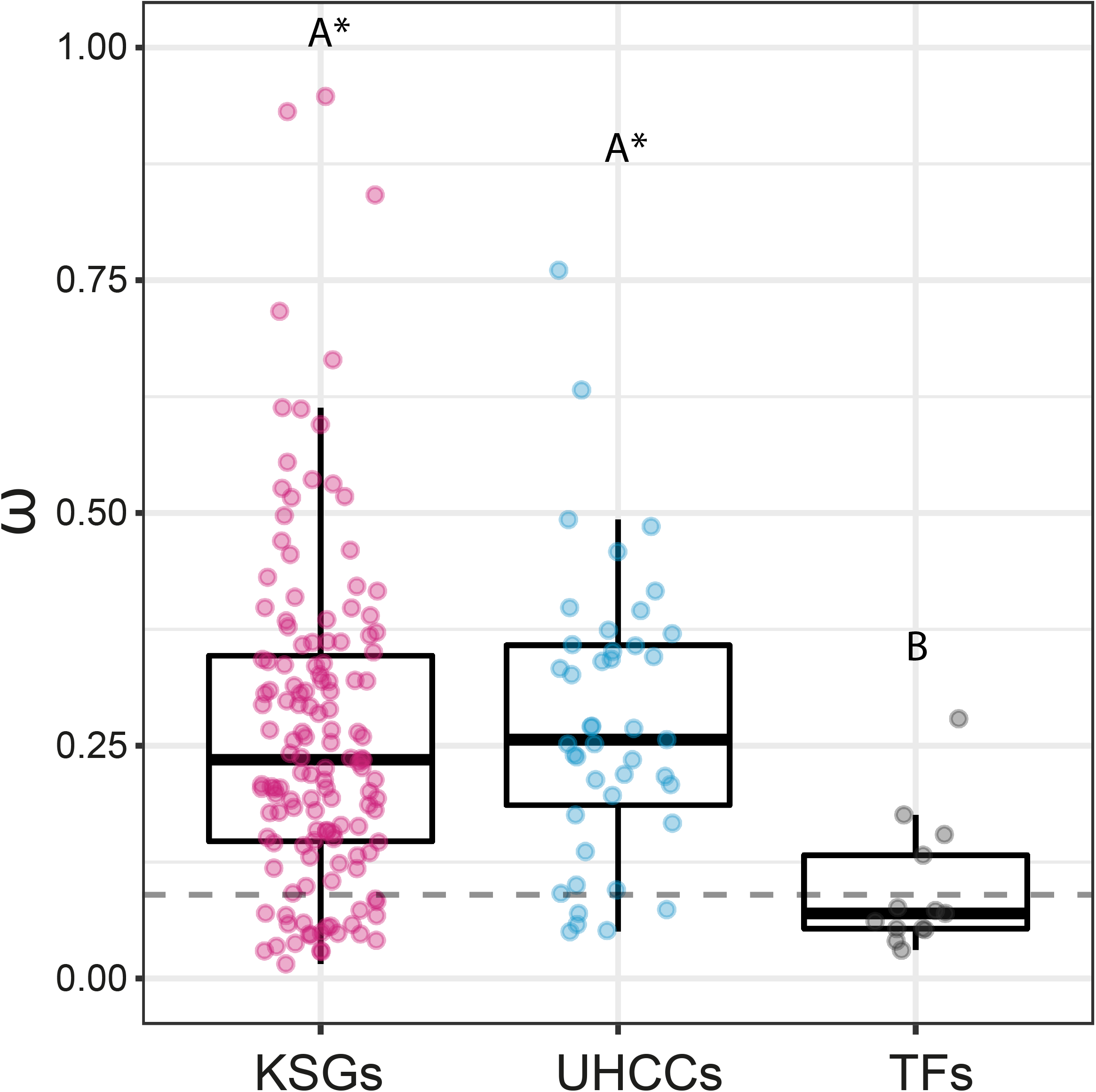
Mean *Ka/Ks* (*ω*) across the *melanogaster* group for Known Seminal Genes (KSGs), Unconfirmed High Confident Candidates (UHCCs), and candidate transcription factors driving the expression of seminal genes in the accessory glands (TFs). TFs searches are described in the Identification section and estimation procedures in Methods (Molecular Evolutionary Analyses). The horizontal discontinuous line represents the mean value for all protein-coding genes [according to Haerty et al. (2007)]. Different letters above boxes indicate differences between groups and * indicates differences between the group and the mean value (GLM followed by Tukey comparisons; p < 0.05).

**Table 2.** Molecular function annotation of Known Seminal Genes (KSGs) and Unconfirmed High Confident Candidates (UHCCs). For each group, count (and percentage) and false discovery rate (FDR) are shown for each GO term found with DAVID with more than one gene.

| GO term | KSGs |  | UHCCs |  |
| --- | --- | --- | --- | --- |
|  | Count | FDR | Count | FDR |
| serine-type endopeptidase inhibitor activity | 18 (10.4%) | 1.76E-16 | 0 | — |
| hormone activity | 6 (3.5%) | 3.09E-04 | 0 | — |
| galactose binding | 5 (2.9%) | 3.09E-04 | 0 | — |
| lipase activity | 5 (2.9%) | 0.00414 | 0 | — |
| serine-type endopeptidase activity | 11 (6.4%) | 0.00804 | 3 (6.5%) | 0.07748 |
| odorant binding | 7 (4.0%) | 0.01045 | 0 | — |
| flavin-linked sulfhydryl oxidase activity | 3 (1.7%) | 0.01045 | 0 | — |
| peptidase inhibitor activity | 3 (1.7%) | 0.01362 | 0 | — |
| carbohydrate binding | 6 (3.5%) | 0.01389 | 0 | — |
| hydrolase activity acting on ester bonds | 4 (2.3%) | 0.02990 | 3 (6.5%) | 0.07739 |
| carboxyesterase activity | 4 (2.3%) | 0.19229 |  |  |
| protein disulfide isomerase activity | 3 (1.7%) | 0.09327 | 0 | — |
| thiol oxidase activity | 2 (1.2%) | 0.18233 | 0 | — |
| unannotated | 77 (44.5%) | — | 37 (80.4%) | — |

Aside from *D. melanogaster*, the only *Drosophila* species in which seminal genes were extensively identified are *D. mojavensis* (Almeida & Desalle 2009; Kelleher et al. 2009; Wagstaff & Begun 2005), *D. pseudoobscura* (Karr et al. 2019), *D. simulans* (Begun & Lindfors 2005; Findlay et al. 2008; Swanson et al. 2001), *D. virilis* (Ahmed-Braimah et al. 2017), and *D. yakuba* (Begun et al. 2006; Findlay et al. 2008). Some (or a few) putative seminal genes were also identified in *D. biarmipes* (Imamura et al. 1998), *D. erecta* (Begun et al. 2006), *D. funebris* (Baumann et al. 1975; Schmidt et al. 1989), *D. mayaguana* (Almeida & Desalle 2009), and *D. suzukii* (Ohashi et al. 1991; Schmidt et al. 1993). Given the good recall of the stringent criteria we used here to identify candidates, we think that other *Drosophila* species could be assayed for seminal genes using similar criteria. Thus, further research on transcriptomic data generated from accessory glands would provide enough starting information to identify at low cost seminal genes in many species.

However, identifying SFPs in multiple species is only part of the equation. The evolution of the seminal proteome may also diverge through changes in the expression level of seminal genes. Begun and Lindfors (2005) found that transcript abundance of the seminal gene *Acp24A4 (FBgn0051779*) differs drastically between *D. melanogaster* and its sibling *D. simulans*. Findlay et al. (2009) reported differences between *D. melanogaster*, *D. simulans*, and *D. yakuba* in the expression level and sex-specificity of several seminal genes. Similarly, Ahmed-Braimah et al. (2017) uncovered large differences in seminal transcripts abundance between members of the *virilis* subgroup. Although these studies documented divergence between closely related species for seminal genes at the regulatory level, neither the cis nor the trans regulatory elements have been studied in depth.

Transcription is a key control point of gene expression, thus the evolution of transcription factors (TFs) that are expressed in the male accessory glands may explain much of the changes in expression of seminal genes across species. However, most of the accessory glands TFs have yet to be identified. To our knowledge, the only known accessory glands’ TFs are the hox gene *Abd-B (FBgn0000015*), the homeodomain transcription repressor *dve (FBgn0020307*), and the paired-rule gene *prd (FBgn0003145*), which are required for the normal development of accessory glands and the production of functional ACPs (Gligorov et al. 2013; Minami et al. 2012; Xue & Noll 2002). Nevertheless, these genes encode pleiotropic master regulators involved in the morphogenesis of several organs and may be subjected to strong evolutionary constraints. Therefore, future research focused on the identification of accessory glands TFs will advance our understanding of how seminal genes’ expression has evolved.

It can be argued that TFs implicated in the regulation of seminal genes’ expression (seminal TFs) correlate with seminal genes in transcript abundance. Ayroles et al. (2011) found 224 *D. melanogaster* genes that, besides being expressed in male reproductive tissues, showed correlated expression patterns to at least seven KSGs. Therefore, we updated this list to the current release (FlyBase r2020_03) and searched it for accessory glands TFs using an online prediction tool implemented in AnimalTFDB3.0, a comprehensive database of animal TFs (Hu et al. 2019). This first search led to the identification of eight putative seminal TFs, including the known *prd* and genes with unknown function (e.g., *FBgn0034870, FBgn0030933*, and *FBgn0028480*). We confirmed that all these candidates are distinctly expressed in the male accessory glands according to both modENCODE (implemented in FlyBase r2020_03) and FlyAtlas2.

Expression pattern does not necessarily correlate between seminal genes and seminal TFs. Thus, we made a second search of TFs in a more extensive list of genes including all those whose expression is enriched in the male accessory glands according to modENCODE (no less than ‘Moderately High’ in accessory glands but no more than ‘Moderate’ in any non-reproductive adult tissues) and FlyAtlas2 (accessory glands enrichment higher than 1). This second search retrieved most of the genes found in the first search plus six new candidates that have not been implicated in reproduction (table 3).

**Table 3.** *D. melanogaster* seminal transcription factors candidates. Aligment e-value and the assigned DNA-binding domain family are shown for each candidate found with AnimalTFDB3.0. The first search was performed on genes whose expression strongly correlates to KSGs expression according to Ayroles et al. (2011). The second search was performed on genes whose expression is enriched in the male accessory glands according to modENCODE and FlyAtlas2 *D. virilis* search, which was performed using Blastp (alignment bit score > 80), shows the presence/absence of homologs among the *D. virilis* putative seminal TFs.

| Name or symbol | FlyBase ID | TF family | e-value | First search | Second search | *D. virilis* search |
| --- | --- | --- | --- | --- | --- | --- |
| retn | FBgn0004795 | ARID | 3.10E-22 | Yes | No | No |
| CG7556 | FBgn0030990 | MYB | 5.00E-16 | Yes | Yes | No |
| prd | FBgn0003145 | PAX | 1.10E-71 | Yes | Yes | Yes |
| toe | FBgn0036285 | PAX | 1.00E-33 | Yes | Yes | Yes |
| CG13559 | FBgn0034870 | zf-LITAF-like | 5.30E-17 | Yes | Yes | No |
| CG6470 | FBgn0030933 | zf-C2H2 | 0.00020 | Yes | Yes | No |
| CG17841 | FBgn0028480 | TRAM_LAG1_CLN8 | 2.60E-63 | Yes | No | No |
| Myc | FBgn0262656 | bHLH | 5.90E-11 | Yes | No | Yes |
| CrebA | FBgn0004396 | TF_bZIP | 3.20E-15 | No | Yes | Yes |
| stc | FBgn0001978 | zf-NF-X1 | 1.10E-10 | No | Yes | Yes |
| CG3065 | FBgn0034946 | zf-H2C2_2 | 4.60E-22 | No | Yes | Yes |
| Bap111 | FBgn0030093 | HMG | 1.30E-16 | No | Yes | No |
| pzg | FBgn0259785 | zf-C2H2 | 7.50E-09 | No | Yes | Yes |
| CG11414 | FBgn0035024 | zf-C2H2 | 7.00E-05 | No | Yes | No |

Next, we explored whether the candidate TFs we identified in *D. melanogaster* are also expressed in the male accessory glands of *D. virilis*, where accessory glands-biased transcripts were recently identified by RNA-seq (Ahmed-Braimah et al. 2017). Seven of the 14 *D. melanogaster* candidates showed clear homology to *D. virilis* genes with accessory glands-biased transcripts that were also predicted to encode TFs (table 3). This contrasts with the low proportion (16.9%) of *D. melanogaster* seminal genes having homologs among *D. virilis* seminal genes. In addition, *Ka/Ks* ratios estimated for the candidate seminal TFs (0.10 on average, range: 0.03–0.28,) were lower than those estimated for seminal genes (0.27 on average, range: 0.02–1.51) (fig. 3). These results suggest that the high turnover rate and the rapidly adaptive evolution of SFPs do not have a strong correlate in the evolution of seminal TFs.

The evolution of seminal genes’ regulatory networks may follow the evolution of cis elements rather than that of TFs. However, enhancers, insulators, and promoters that are active in the male accessory glands have not been thoroughly investigated. Thus, the study of seminal TFs and their binding sites is an important area for future research.

Besides TFs and their binding sites, post-transcriptional factors such as microRNAs (miRNAs) are also involved in the regulation of seminal genes’ expression. Recently, Mohorianu et al. (2018) made an important contribution to the understanding of seminal regulatory networks by assessing the role of miRNAs in the modulation of ejaculate composition. The authors found evidence for the presence of several regulatory miRNAs that bind to a given sequence of the 3’ untranslated region (UTR) of seminal transcripts, likely repressing translation. Each miRNA targets a specific group of seminal genes that share the corresponding 3’ UTR target site, which provides males with a mechanism to adjust ejaculate composition (Mohorianu et al. 2018). These findings indicate that seminal genes UTRs and accessory glands miRNAs may have been involved in the evolution of the seminal proteome.

Beyond the regulatory elements identified in *D. melanogaster*, causes underlying the divergence of seminal genes at the regulatory level remain mostly unknown. Certainly, comparative genomics will help to address this problem, however, we first need to identify the involved elements in other species. Therefore, future research studying accessory glands transcriptome in different *Drosophila* species will likely benefit this unexplored field.

## Turnover Rate

One of the most striking characteristics of SFPs is their rich diversity, which seems to be causally related, at least in part, to sexual conflict (Chapman 2008, 2018). In theory, postmating sexual selection can escalate the evolutionary tension between the fitness interests of males and females because male adaptations to sperm competition can be harmful to females (Chapman et al. 1995; Lung et al. 2002; Mueller et al. 2007). Selection will then favor both female traits that counteract detrimental male adaptations and male traits that respond to female resistance, potentially leading to coevolutionary arms races between male persistence and female resistance (Arnqvist 2004; Chapman et al. 2003). SFPs, by affecting female physiology and behavior, clearly influence fertilization success and sperm competitiveness. Therefore, sexual antagonistic coevolution between SFPs and the female counterparts likely accounts for the rapid divergence of seminal proteomes (Sirot et al. 2014).

As sperm competition and sexual conflict can lead to rapid adaptive divergence of orthologous SFPs, they may also promote divergence of the seminal protein repertoire through the gain of novel seminal genes as well as through seminal gene loss. On one hand, females will not be adapted to resist the action of novel SFPs. On the other hand, the expression of ancient SFPs— whose action has been neutralized by females’ counter-adaptations—will not be sustained by natural selection. According to this hypothesis, turnover of seminal genes would be adaptive for males because it would provide males with resources to “stay ahead” of female resistance (Chapman 2018; Sirot et al. 2014). Evidence supporting sexual conflict as a driver of seminal protein evolution abounds and comes from diverse sources (reviewed in Chapman 2018; Hollis et al. 2019; Sirot et al. 2014).

High turnover rate of seminal gene sets was first noted by Wagstaff and Begun (2005). Assaying the just released *D. pseudoobscura* genome for orthologs of *D. melanogaster* ACP-coding genes, the authors noticed an unexpectedly high proportion of absences, suggesting that an important number of seminal genes are lineage-specific. Later that year, Begun and Lindfors explored the presence/absence patterns of three *D. simulans* ACP-coding genes across closely related species of the *melanogaster* subgroup, to which *D. simulans* belongs. They found that two of these genes *(Acp23D4* and *Acp54A1*) were absent in at least one species but had one to three copies in the rest. Mueller et al. (2005), by performing comparative sequence analysis on 52 ACP-coding genes of the *melanogaster* subgroup, found that 22 of them were not conserved in *D. pseudoobscura*. Overall, these studies introduced the idea that the fraction of the genome encoding SFPs is, by means of gene gain and loss, unusually dynamic.

With the release of the genomes of 12 *Drosophila* species (Drosophila 12 Genomes Consortium 2007), several comparative studies confirmed this pattern (e.g., Ahmed-Braimah et al. 2017; Findlay et al. 2008, 2009; Haerty et al. 2007; Zhang et al. 2007). However, since too few dipteran species were assayed for extensive identification of seminal genes, a comprehensive analysis to trace the origin and loss of seminal genes in a phylogenetic context is lacking. Currently, we do not know, for instance, to what extent orthologs of *D. melanogaster* seminal genes also encode SFPs in other species of the genus. We do not know either how long ago these genes have encoded SFPs in the *D. melanogaster* lineage. Identifying seminal genes/proteins in other *Drosophila* species would allow to not only survey the evolutionary history of SFPs but also study how new SFPs arise and how regulatory elements of seminal genes diverge between species. So far, these questions have been barely explored.

Another question that arises is whether a core of SFPs playing essential reproductive roles has been conserved throughout evolution. In such a case, these “essential SFPs”, critical for reproduction, should be present in a vast number of taxa. They could be searched by recognizing the SFPs shared not only by closely related species but also by several phylogenetically distant taxa; those shared only by closely related species would include both essential and non-essential ones.

Intending to survey this hypothesis in Diptera, here we compiled a list of SFPs of the *melanogaster* subgroup (those identified in *D. melanogaster, D. simulans* and/or *D. yakuba*) and search it for homologs among SFPs identified in other dipteran taxa with well-known seminal genes/proteins [see methodological procedures in Methods (Orthology of SFPs among Diptera)].

Taking into account that identification studies are hardly exhaustive, we only considered the three outgroup taxa for which SFPs or seminal genes were identified in no less than two species by independent extensive searches. These taxa were the *virilis-repleta* radiation of the *Drosophila* subgenus (that split from *D. melanogaster* ~35 mya), tephritid fruit flies (that split from *D. melanogaster* ~120 mya), and mosquitoes (that split from *D. melanogaster* ~250 mya). We clustered all annotated proteins of 19 *Drosophila* species, including the *melanogaster* subgroup and the *virilis-repleta* radiation, in 23782 groups of orthologs (orthogroups), 196 of which have at least one seminal gene of the *melanogaster* subgroup. Among these 196 orthogroups 41 contain seminal genes of the *virilis-repleta* radiation, 11 have at least one homolog of tephritid seminal genes, and 25 have at least one homolog of mosquitoes’ seminal genes (fig. 4). Caution should be taken when comparing these numbers because they relied on different homology criteria, some applied by different previous studies (supplementary table S2). However, considering that 11,298 out of the 13,969 (81%) protein-coding genes have certainly clear homologs in mosquitoes (blastp bit score > 50), the number of SFPs shared by the four evaluated taxa seemed to be remarkably low: only two orthogroups had seminal genes of the four taxa. One of these orthogroups contains only one *D. melanogaster* seminal gene *(FBgn0034753*), which encodes a peptidyl-prolyl cis-trans isomerase. The other contains five *D. melanogaster* seminal paralogs that encode protease inhibitors with Kazal domains and belong to a tandem gene cluster located in the left arm of the second chromosome. Within this gene family, we found *FBgn0266364*, which was identified as a novel candidate in the present report, and *FBgn0051704*, which is reported in FlyBase r2020_03 as ortholog of *SPINK2*, a human gene implicated in male infertility.

**Fig. 4.**
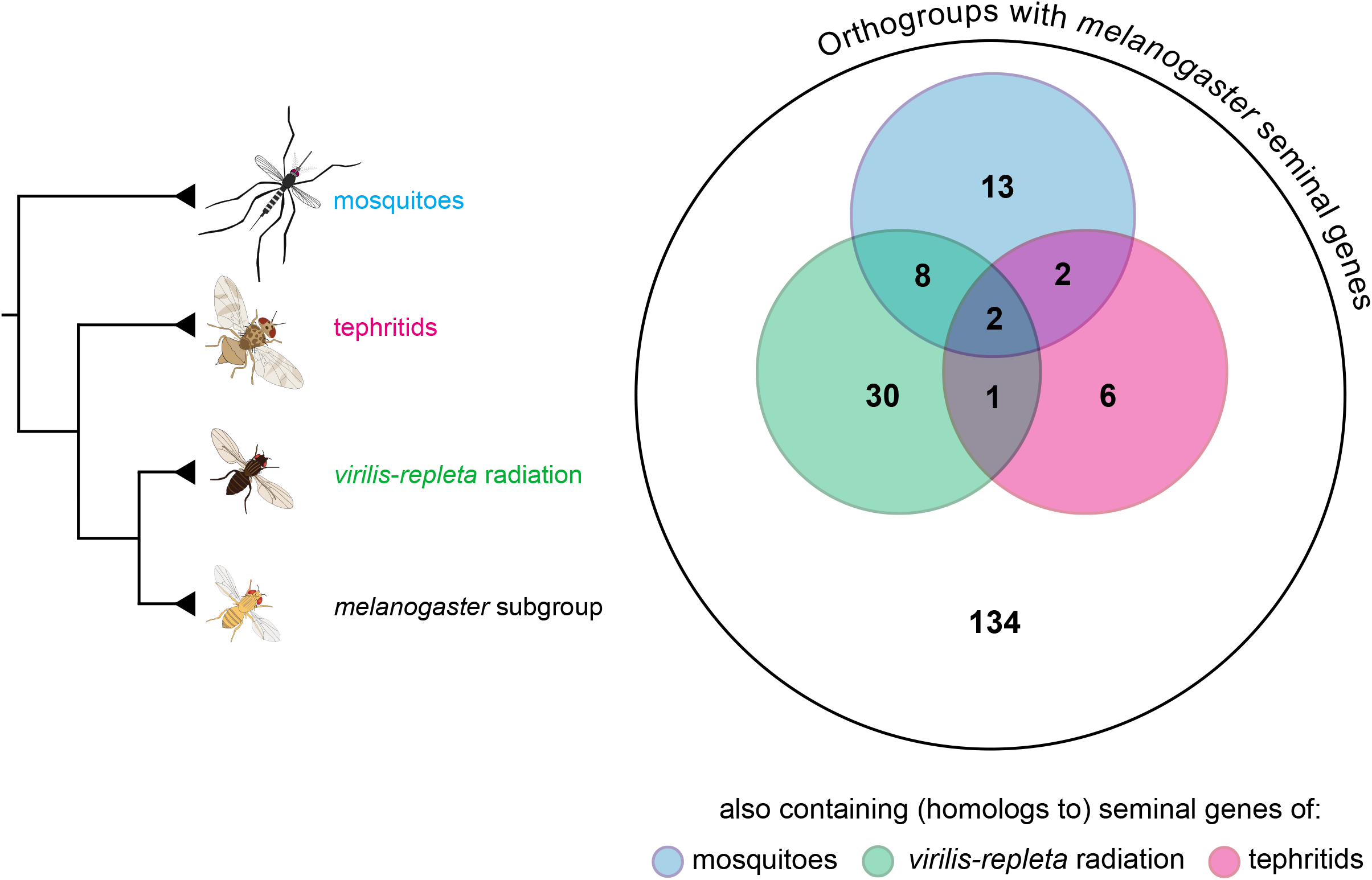
Seminal genes shared between the *melanogaster* subgroup and other Diptera. Numbers refer to the 196 *Drosophila* orthogroups (generated with Orthofinder) having at least one seminal gene of the *melanogaster* subgroup. Orthogroups having seminal genes of various taxa are represented by overlapped areas.

Although the number of taxa included in our analysis is low, the results indicate that most SFPs in Diptera are lineage-specific, which strongly suggests that most SFPs have a short evolutionary life (or diverges rapidly beyond detectable homology) and that not many—if any—have been critical for reproduction throughout Diptera evolution. Still, even if the seminal protein repertoires of the taxa we analyzed were fairly complete, our results would be far from being conclusive because homology detection across dipteran families can be inefficient for rapidly evolving seminal genes. In this sense, it would be more feasible to search for “essential SFPs” within specific groups of the *Drosophila* genus. However, the repertoire of SFPs is currently known for too few species. Thus, the search for “essential SFPs” within *Drosophila* must await more studies assaying SFPs in a wider spectrum of species.

Despite those observations and claims, gene birth and death rates were never estimated for SFPs. To obtain these estimates, we pruned the 196 orthogroups containing *D. melanogaster* seminal genes, leaving only the nine species for which genomic annotations were updated at least once [see Methods (Gene Birth and Death Rates)]. Then, duplications, losses, and orthogroup gains were identified in the gene trees of each orthogroup (fig. 5) and each event rate was estimated from the obtained figures. Taking into account divergence dates reported in Obbard et al. (2012), the estimated duplication rate was 0.0097 duplications per gene per million years (/gene/my) and the loss rate was 0.0122 losses/gene/my (0.0133 duplications/gene/my and 0.0212 losses/gene/my considering only the species of the *melanogaster* group). The species with the greatest gene loss rate was *D. sechellia* (49 losses), which could be an artifact of genome sequencing, assembly, and annotation. However, the number of protein-coding genes annotated for this species is the highest in the *melanogaster* group and a similar pattern of high gene loss was previously observed for olfactory genes in this species (Almeida et al. 2014; McBride 2007). The authors associated this with *D. sechellia* specialization and endemism, which could also have implications for the mating system and reproductive proteins. Regarding orthogroup gains in the *D. melanogaster* lineage, the estimated rate was 0.0047 gains/gene/my and the total number of identified events was 87. The acquisitions were inferred in the ancestors of the *Sophophora* subgenus (25), the *melanogaster* group (22), the *melanogaster* subgroup (35), and the *melanogaster* complex (*D. melanogaster, D. simulans*, and *D. sechellia*) (5). Interestingly, the latter figure accounts for more than half the number of putative *de novo* genes identified by Zhou et al. (2008) in the *melanogaster* complex.

**Fig. 5.**
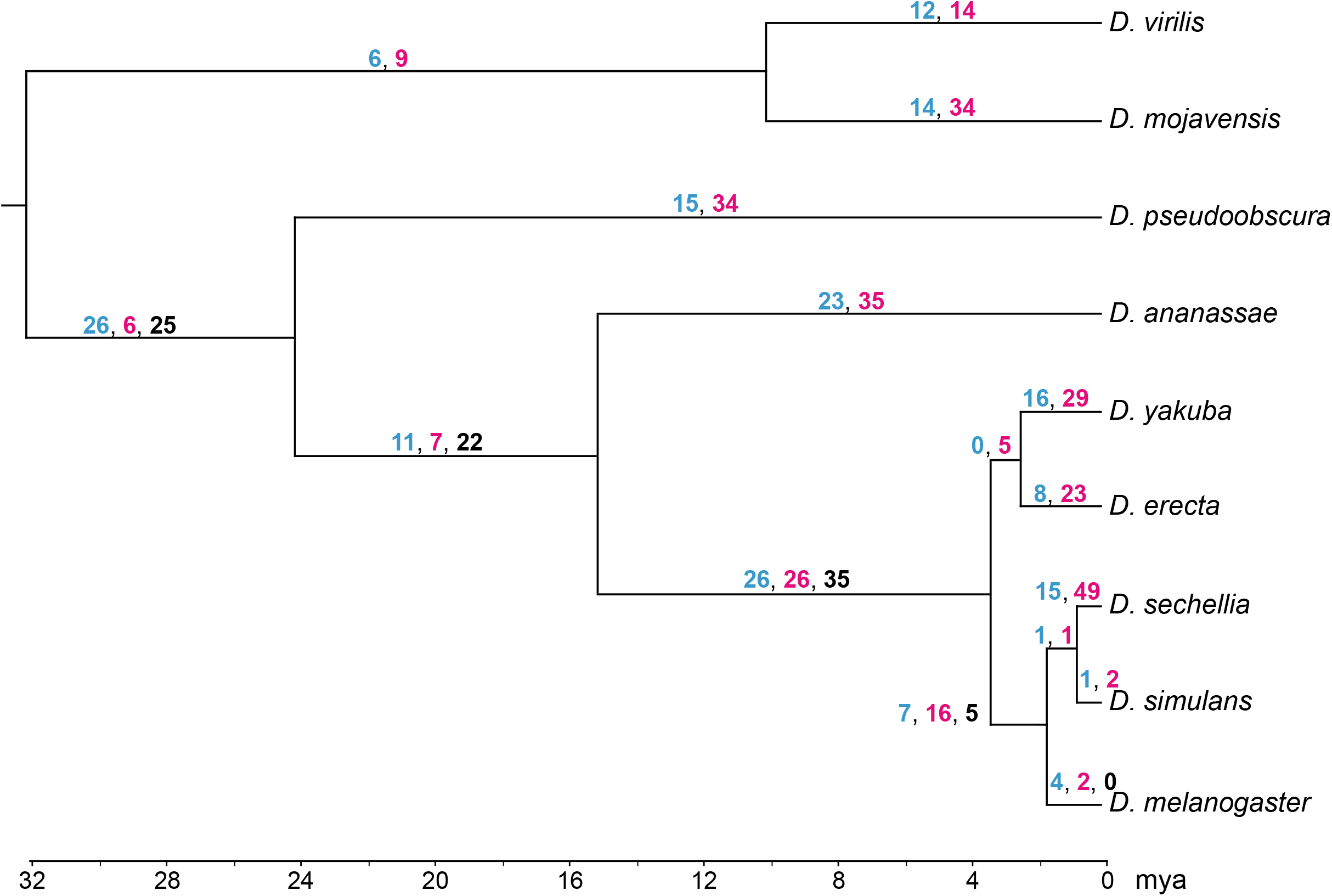
Duplication (blue), loss (magenta), and *de novo* emergence (black) events among orthogroups containing *D. melanogaster* seminal genes. The numbers of events are shown per branch. Since orthogroups without *D. melanogaster* SFPs were not considered, *de novo* gains for branches outside the *D. melanogaster* lineage, which are zero, are not shown. Divergence times were obtained from Obbard et al. (2012).

Using 12 *Drosophila* genomes, Hahn et al. (2007) estimated a total event (gene duplications + losses) rate of 0.0013 events/gene/my based on Tamura et al. (2004) divergence dates. Using the same dates, we estimated for the *D. melanogaster* seminal genes an event rate of 0.0096 events/gene/my (0.0111 events/gene/my considering only the species of the *melanogaster* group). This suggests that seminal genes’ families, though they may not contain seminal genes of non-*D. melanogaster* species, are approximately seven times more dynamic than the average gene family in *Drosophila*.

## Mechanisms of Origin

The high turnover rate in seminal genes/proteins repertoires implies a high proportion of novel seminal genes/proteins restricted to young lineages or unique species. This facilitates studying the evolution of novel genes in a common cellular background (i.e., accessory glands) in groups of closely related species, where the molecular routes of gene origin are more likely traceable. Thus, seminal genes provide an excellent opportunity to investigate how novel proteins and biological functions emerge. Four mechanisms have been reported or proposed so far as responsible for the origin of seminal genes in *Drosophila*: duplication of seminal genes, duplication of non-seminal genes, gene co-option into the male reproductive tract, and *de novo* evolution (reviewed in Sirot 2019).

The first mechanism proposed was duplication of preexisting seminal genes (e.g., Almeida & Desalle 2009; Findlay et al. 2008; Holloway & Begun 2004; Mueller et al. 2005; Wagstaff & Begun 2005). When a seminal gene is entirely duplicated so that both copies, the new and the old, encode the same SFP, ensuing mutations may lead to subfunctionalization or neofunctionalization, giving rise to novel SFPs with similar amino acid sequences. Most of the seminal genes encoding these proteins are located in clusters of nearby genes on the second chromosome (fig. 2), showing that tandem duplication followed by mutation has played an important role in the divergence of the seminal proteome. For instance, *FBgn0043825*, *FBgn0051872*, and *FBgn0265264* are three paralogs located in tandem on the left arm of the second chromosome, which encode SFPs with triglyceride lipase activity (Mueller et al. 2005).

Duplication of genes that are not expressed in the male reproductive system and do not encode SFPs may also be a source of novel seminal genes (Sirot 2019); if a duplicate ends up placed under the control of regulatory elements driving its expression in the accessory glands, it may become a new seminal gene. Genes encoding proteins that already have secretion signals are likely sources for this mechanism. An example of this is the origin of the seminal gene *FBgn0052833*, which resulted from a duplication-mediated co-option of a female-expressed gene whose original copy encodes a secretory protein of the sperm storage organs (Sirot et al. 2014). Another example comes from odorant binding proteins (OBPs), a highly dynamic family of olfactory genes that are usually expressed in the antennae. Four OBP genes, however, have been co-opted into the accessory glands exclusively in the lineage leading to the *melanogaster* group (Almeida et al. 2014). Interestingly, the rates of protein evolution of these genes were the highest among OBPs.

Although duplication may facilitate sequence or expression evolution because of initial redundancy (one copy can change, while the other maintains the original function), some *Drosophila* seminal genes seem to have arisen via gene co-option in the absence of a previous gene duplication event (Findlay et al. 2008). *FBgn0262571*, a *D. melanogaster* seminal gene exclusively expressed in the male accessory glands, belongs to a single-copy gene family (Sepil et al. 2019). Its orthologs, despite encoding proteins with secretion signal, are not within the repertoire of seminal genes in either *D. mojavensis, D. pseudoobscura*, or *D. virilis* (the only non-*melanogaster* group species of the genus in which seminal genes were extensively identified). Therefore, despite not being duplicated, this gene was potentially co-opted into the accessory glands in the *D. melanogaster* lineage, during the evolution of the *melanogaster* group.

Some other seminal genes may have emerged *de novo* from ancestrally noncoding DNA (Begun et al. 2006; Findlay et al. 2008; Haerty et al. 2007). While sperm competition and sexual conflict may steadily select for innovation in the male ejaculate, “fitness valleys” limit the paths available for the evolution of preexisting proteins (Camps et al. 2007). In this sense, young *de novo* seminal genes may be less constrained and may have more opportunities to fill the emerging functional niches. Curiously, the first evidence consistent with *de novo* gene birth comes from studies aimed to identify genes specifically expressed in *Drosophila* male accessory glands (Begun et al. 2006) or testes (Begun et al. 2007; Zhao et al. 2014). Given the high proportion of insect seminal genes without identified orthologs, *de novo* gene birth is believed to account for the origin of many seminal genes (reviewed in Sirot 2019). So far, however, no *Drosophila* seminal genes have yet been identified as *de novo* genes with high confidence, possibly because distinguishing *de novo* birth from horizontal transfer or rapid protein divergence (which is common among seminal proteins) is challenging (Zile et al. 2020).

Despite particular cases, a broad-scale analysis to determine the relative contribution of the alternative mechanisms of origin has yet to be completed. In an attempt to discern which of the mentioned mechanisms were responsible for the origin of young *D. melanogaster* SFPs [those that have arisen during the evolution of the *melanogaster* species group, i.e., less than ~25 million years ago (mya)], we identified gene families that included *melanogaster* group’s seminal genes. Given that homology detection power banishes with divergence, evaluating alternative mechanisms of origin for older genes would be much more uncertain. Gene families were obtained by clustering the proteins of reference proteomes of 19 *Drosophila* species [see Methods (Seminal Gene Families)]. This analysis revealed that our set of 219 *D. melanogaster* SFPs belong to 168 gene families. To determine which seminal genes have likely emerged after the origin of the *melanogaster* group (which were dubbed young seminal genes), and to infer the most likely mechanism of origin, we manually inspected the gene family tree of all these 168 gene families. Specifically, we explored the presence/absence of orthologs and paralogs, and whether they had been classified as SFPs. We then applied the parsimony principle to determine, according to the observed pattern, which mechanism was most likely responsible for the origin of each young *D. melanogaster* SFP (fig. 6 illustrates our criteria). See Methods (Seminal Gene Families) for a more detailed description of the applied criteria. In cases where *n* mechanisms were equally likely, we assigned *“1/n* genes” to each mechanism.

**Fig. 6.**
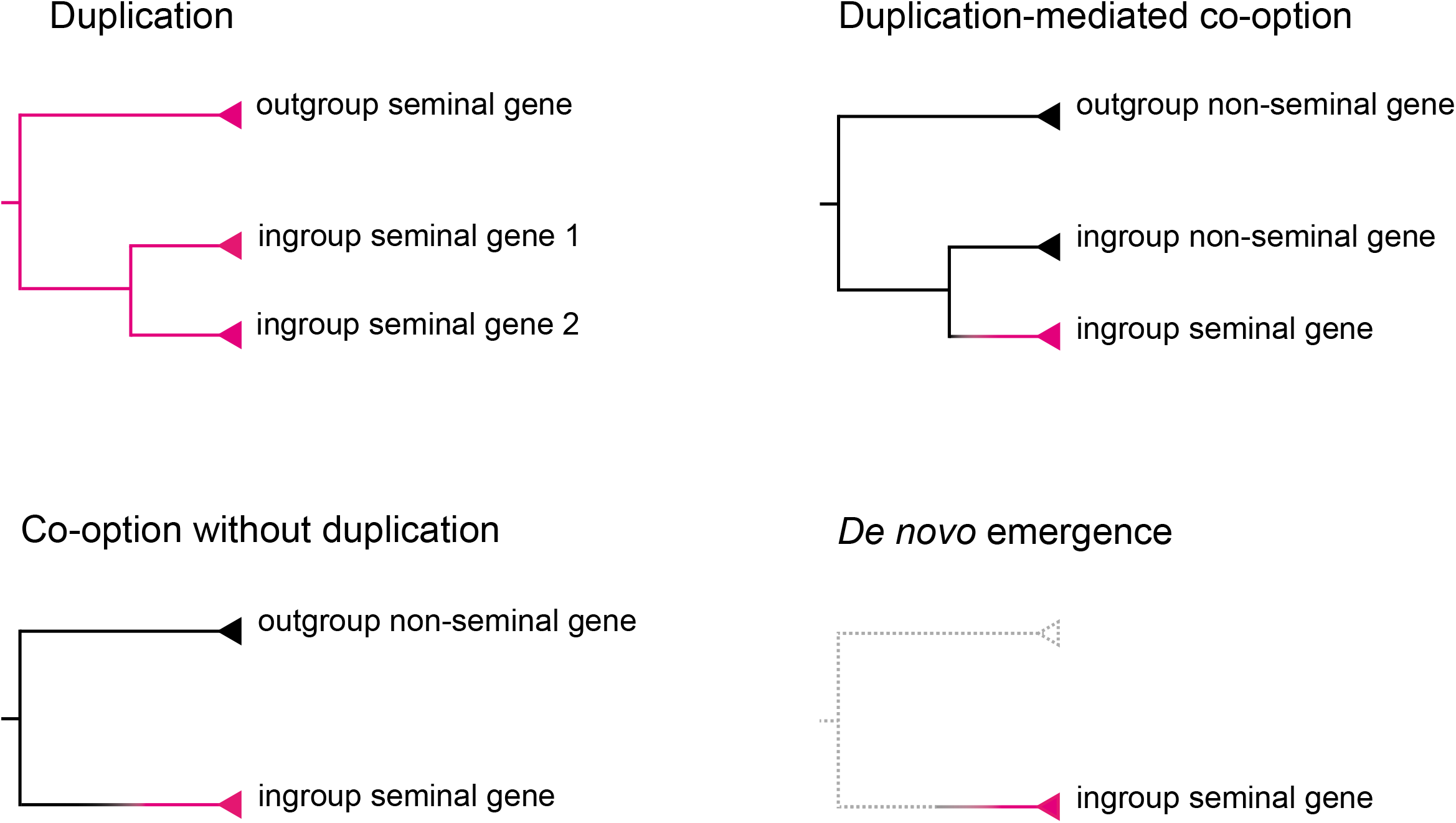
Expected gene family topology for each considered mechanism of seminal gene origin. Ingroup genes represent *melanogaster* genes, while outgroup genes represent genes of any non-*melanogaster* group species for which seminal genes are known. Magenta branches correspond to seminal genes, while black branches correspond to non-seminal genes. Grey discontinuous branches stand for the absent of homologs.

In this way, we estimated that 76 *D. melanogaster* seminal genes existed as seminal genes (before the split from the lineage leading to *D. pseudoobscura* (~25 mya). For 13 seminal genes, we could not determine whether the origin was before or after that split since they exhibited uncertain homology to sequences of outgroup or distant species. Among the remaining 130 *D. melanogaster* seminal genes (i.e., the tentatively young ones), we classified ~27 (20.6%) as duplicates of preexisting seminal genes, ~7 (5.3%) as co-opted duplicates (duplicates of non-seminal genes), ~47 (36.5%) as co-opted without duplication, and ~49 (37.6%) as putative orphans (fig. 7).

**Fig. 7.**
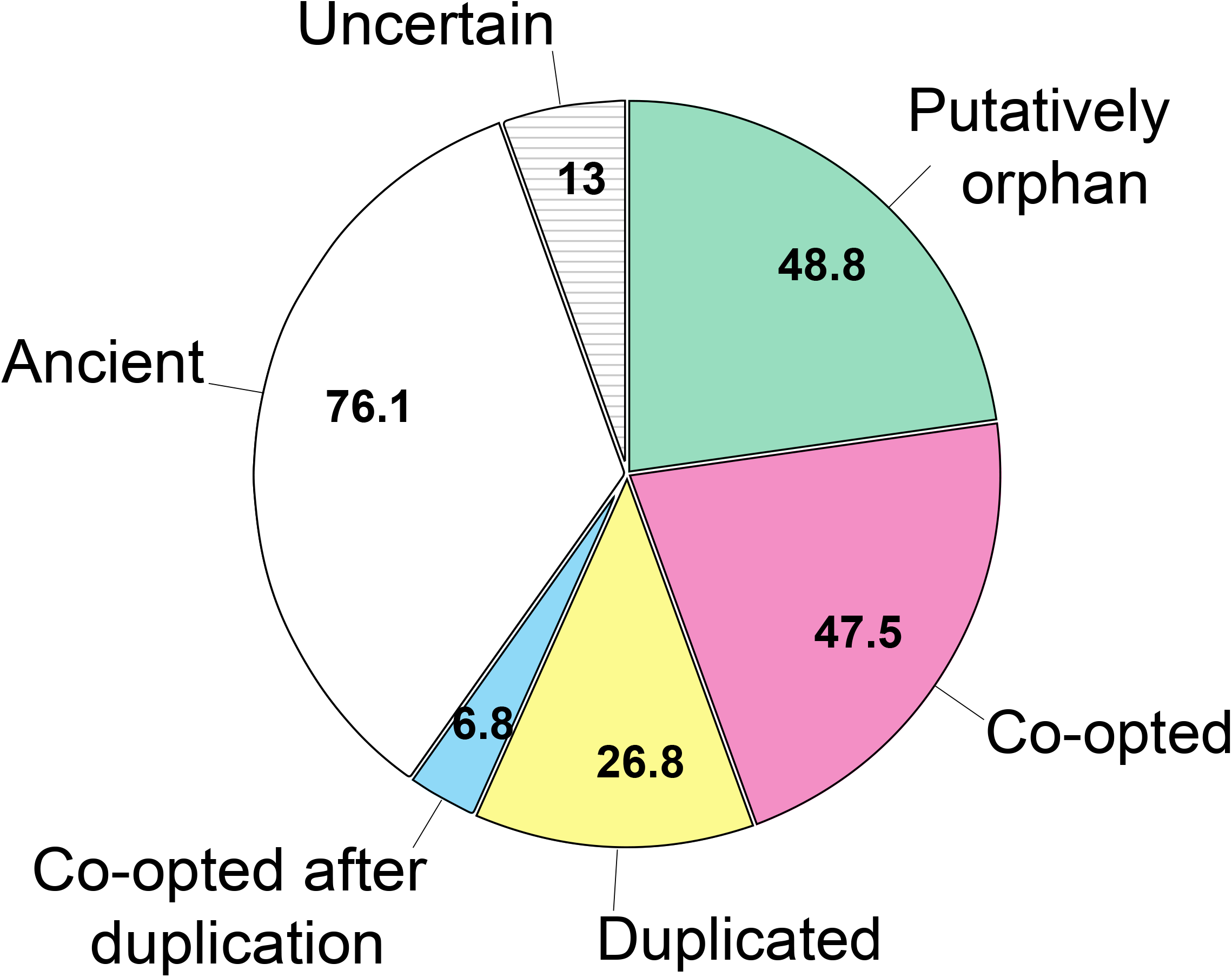
Most likely mechanisms of origin of *D. melanogaster* seminal genes. Mechanisms were proposed according to our analysis of seminal gene families only for tentatively young seminal genes, i.e., those that have likely emerged after the split from the lineage leading to *D. pseudobscura*. Uncertain genes represent those we could not determine whether they are young or ancient.

These results may give the impression that *de novo* emergence was responsible for the origin of many *D. melanogaster* seminal genes. However, our approach did not contemplate all possible mechanisms of gene origin and may have confounded some. For instance, a non-orphan seminal gene showing fast evolution may have diverged beyond detectable homology and be construed as an orphan gene. Some of the proteomes we used may be incomplete due to potentially defective genomic annotations, which may also have led to the overestimation of taxonomically restricted genes. In consequence, the actual number of orphans among seminal genes of the *melanogaster* group is surely lower than the one we estimated. In fact, we could not ensure *de novo* status for any of the identified putative orphans [see applied criteria in Methods (*De Novo* Status Validation)]. Briefly, after examining several *Drosophila* annotated genomes, we failed to find taxonomically restricted seminal genes with syntenic homologous, reliably noncoding sequences in any outgroup species. This means that these gene families, which were initially identified as taxonomically restricted to the *melanogaster* group, may be classified as originating through rapid evolution (among other mechanisms) rather than *de novo* emergence. Therefore, the relative contribution of *de novo* emergence to the origin of *Drosophila* seminal genes may be more limited than previously thought. Gene co-option, on the other hand, appears to be the most frequent mechanism of origin.

To uncover the possible ancestral expression pattern of those few seminal genes that, according to our analysis, appear to have arisen via duplication-mediated co-option, we checked the expression pattern of the respective non-seminal paralogs. According to modENCODE (implemented in FlyBase r2020_03), these paralogs are expressed in the larval salivary gland, the adult female spermatheca, the pupal fat body, or the adult digestive system. Whether these tissues represent common sources for co-option into the seminal fluid will require further cross-species exploration of co-opted seminal genes (for examples in other insects see Martinson et al. 2017; Meslin et al. 2015).

Alternative mechanisms of seminal genes’ origin—such as exon/domain shuffling, gene fission/fusion, horizontal gene transfer, and reading-frame shift—should be explored in the future. Also, further identification of SFPs in more *Drosophila* species will allow for more accurate discrimination between alternative mechanisms, for dating gene origin more precisely, and for exploring gene origin in other groups.

## Conclusions

Here, we provided an overview of the inter-specific divergence of *Drosophila* SFPs summarizing the current state of knowledge and emphasizing the intriguing aspects that are less understood. We focused on the conservation of SFPs across the order Diptera and the mechanisms of origin of *Drosophila* seminal genes. We not only inspected some of the main contributions to these topics but also compiled genomic information from multiple species and performed molecular evolutionary analyses to address some broad questions that remain open.

Using reviewed criteria, we presented a novel set of high-confidence seminal protein candidates for *D. melanogaster* and generated a database of *Drosophila* SFPs. We also provided, for the first time, a list of accessory glands (putative or confirmed) TFs presumptively controlling the expression of SFPs.

Two interesting patterns derive from our comparative genomic analyses. First, given the low number of common SFPs found among the three inspected dipteran families, the hypothesis that there is a core of indispensable, “essential SFPs” conserved across Diptera seems unlikely. Second, gene co-option appears to be the most frequent mechanism accounting for the origin of *Drosophila* seminal genes. As *de novo* evolution could not be ensured for any seminal gene, our analysis failed to support the hypothesis that *de novo* emergence is a frequent mechanism of origin for seminal genes.

Despite the insights we have gained, it is evident that characterizing the seminal proteome in more species, especially in those outside the *melanogaster* group, is imperative to fill important knowledge gaps. While proteomics on isotopic labeled flies and quantitative proteomics have proven to be useful to carry out this task, our searches suggest that RNA-seq on accessory glands, which is less challenging and cheaper, would provide valuable starting information.

## Methods

### Orthology of SFPs among Diptera

Supplementary table S2 summarizes the sources of the list of SFPs for each considered taxa (lists are available upon request). To identify the orthologs of the SFPs identified in the *melanogaster* group (ingroup), we employed the following strategy. First, we gathered the proteomes of 19 *Drosophila* species (see below) and used Orthofinder, a platform for comparative genomics (Emms & Kelly 2015, 2019), to cluster the proteins in groups of orthologs (orthogroups). Then, we searched for the orthogroups that had any SFP of the *melanogaster* subgroup [i.e., the 219 of *D. melanogaster* or those of *D. simulans* and/or *D. yakuba* identified by Findlay et al. (2008)]. The input protein sequences were obtained from reference proteomes available in FlyBase, NCBI, or specific genome projects’ sites. The *Drosophila* species of the *melanogaster* group included in the analysis were *D. ananassae* [dana_r1.06 (FlyBase r2020_03)], *D. biarmipes* [Dbia_2.0 (Richards et al. unpublished, NCBI)], *D. bipectinata* [Dbip_2.0 (Richards et al. unpublished, NCBI)], *D. elegans* [Dele_2.0 (Richards et al. unpublished, NCBI)], *D. erecta* [dere_r1.05 (FlyBase r2020_03)], *D. eugracilis* [Deug_2.0 (Richards et al. unpublished, NCBI)], *D. ficusphila* [Dfic_2.0 (Richards et al. unpublished, NCBI)], *D. kikkawai* [Dkik_2.0 (Richards et al. unpublished, NCBI)], *D. mauritiana* [dmauMS17_r1.0 (Nolte et al. 2013)], *D. melanogaster* [dmel_r6.34 (FlyBase r2020_03)], *D. rhopaloa* [Drho_2.0 (Richards et al. unpublished, NCBI)], *D. sechellia* [dsec_r1.3 (FlyBase r2020_03)], *D. simulans* [dsim_r2.02 (FlyBase r2020_03)], *D. suzukii* (Joanna C. Chiu 2020, personal communication), *D. takahashii* [Dtak_2.0 (Richards et al. unpublished, NCBI)], and *D. yakuba* [dyak_r1.05 (FlyBase 2017_03) re-annotated by Yang et al. (2018)]. Species belonging to other species groups (outgroups) were *D. mojavensis* [dmoj_r1.04 (FlyBase r2017_03) re-annotated by Yang et al. (2018)], *D. pseudoobscura* [UCI_Dpse_MV25 (Liao et al. unpublished, NCBI)], and *D. virilis* [dvir_r1.06 (FlyBase 2017_03) re-annotated by Yang et al. (2018)]. These three species were chosen because they were the only ones outside the *melanogaster* group in which seminal genes were extensively studied. As Yang et al. (2018) did not annotate CDSs, we predicted for *D. mojavensis, D. virilis*, and *D. yakuba* one protein per gene with RefProt pipeline (Revale & Hurtado, available upon request), which is based on TransDecoder (Haas et al. 2013), Blast (Altschul et al. 1990), HMMER (hmmer.org), and several inhouse R scripts (R-project.org). In our experience, Orthofinder has limited recall when clustering sequences of very distantly related species. Therefore, to recognize orthogroups with SFPs of species outside *Drosophila (Aedes aegypti, Aedes albopictus, Anopheles gambiae, Bactrocera dorsalis*, and *Ceratitis capitata*) we relied on previous orthology assignments based on Blast (supplementary table S2). We considered a SFP to be shared between *melanogaster* subgroup and any given outgroup if the protein was clustered together with an outgroup SFP in the same orthogroup.

### Molecular Evolutionary Analyses

Estimates of the ratio between the rate of non-synonymous substitution (*Ka*) and the rate of synonymous substitutions (*Ks*) can be used as a proxy to investigate the evolutionary forces that shape the evolution of proteins. Close to zero ratios are associated with purifying selection, whereas ratios close or higher than one mean that the gene evolves under neutrality or that some codons are positively selected. We employed PAML-4.8 (Yang 2007) to obtain *w*, a likelihood-based estimator of *Ka*/*Ks*, for each orthogroup.

### Gene Birth and Death Rates

We pruned the 196 orthogroups containing *D. melanogaster* SFP-coding genes (see above) to include only those species with updated genome annotations, leaving in this way the orthologs of *D. melanogaster*, *D. simulans*, *D. sechellia*, *D. erecta*, *D. yakuba*, *D. ananassae*, *D. pseudoobscura, D. mojavensis*, and *D. virilis*. Then we employed the program Notung-2.9.1.5 (Chen et al. 2000; Darby et al. 2017) to identify gene duplications, losses, and *de novo* gains in each orthogroup by comparing gene trees with the species tree. To be conservative and avoid overestimation, we edited the Notung results to remove duplications and losses when there was an even number of genes per species. With the total number of each of these events for each branch of the *Drosophila* phylogeny, we estimated per gene rates by dividing the number of events by the number of genes in the ancestral branches. These events were summed across all branches and the sum was divided by the total phylogeny time to obtain the rates using the formulas described in Vieira et al. (2007). A gene gain was identified for each orthogroup exclusive of a monophyletic clade.

### Seminal Gene Families

Since Orthofinder inference relies on reciprocal best alignment hits, some paralogous sequences ended up grouped in separate orthogroups. Thus, with the aim of identifying paralogous orthogroups, we compared *D. melanogaster* sequences clustered in different orthogroups using Blastp. We then merged orthogroups with aligned sequences into more inclusive gene families. Since we used a conservative bit score cutoff of 80 for filtering hits, the number of recognized gene families probably represent an upper bound of the actual number. Our objective was to determine the origin of *D. melanogaster* seminal genes that had emerged during the evolution of the *melanogaster* group (i.e., after the split from the lineage leading to *D. pseudoobscura*), so we considered the species belonging to other groups as outgroups. We then used the gene trees generated by Orthofinder to investigate the origins of the *melanogaster* group SFPs. Within each orthogroup, the last common ancestor gene between an outgroup seminal gene and a *D. melanogaster* seminal gene was considered as a seminal gene. Similarly, the last common ancestor gene at the root of any orthogroup containing homologs to seminal genes of tephritids or mosquitoes was also considered as a seminal gene. With these considerations, we inferred the most likely mechanism of origin of each *D. melanogaster* seminal gene by manually inspecting the respective gene family tree. Specifically, we explored the presence/absence of orthologs and paralogs among species of the *melanogaster* group and outgroups applying the parsimony principle over gene gain/loss events (fig. 6). In this way, we first distinguished between “ancient” (those that had emerged before the split from the lineage leading to *D. pseudoobscura*, ~25 mya) and tentatively young (those lacking homologs among outgroup seminal genes, that have likely emerged after the split from the lineage leading to *D. pseudoobscura) D. melanogaster* seminal genes. Then, we classified tentatively young seminal genes into the following four categories: duplicated, co-opted after being duplicated, co-opted without being duplicated, and orphan. In those cases where *n* mechanisms were equally likely, we assigned *“1/n* genes” to each mechanism. Some *D. melanogaster* proteins may have evolved very rapidly, hindering homology detection. Thus, in the case of SFPs classified as orphan with our approach, we evaluated distant homology by comparing *D. melanogaster* SFPs to non-redundant proteins sequences from NCBI databases using Blastp (blast.ncbi.nlm.nih.gov). In this case, we admitted hits (bit score > 39) against sequences of any Diptera: those with any bit score higher than 50 were considered to reflect homology while those with bit scores between 39 and 50 were considered uncertain. Also, for each apparent orphan seminal gene, we checked manually the absence of syntenic open reading frames encoding similar proteins (Blastp: bit score > 39 or positives > 60%) in the *D. pseudobscura* genome by using the Ensembl Metazoa genome browser (Howe et al. 2019).

### *De Novo* Status Validation

To validate the *de novo* status of the putative orphans, we used the conservative criteria applied by Zile et al. (2020). Briefly, as *de novo* genes should have syntenic, homologous noncoding sequences in closely related outgroup species, we inspected each orphan candidate for syntenic, homologous noncoding sequences in well-annotated genomes of outgroup species. Particularly, we examined the latest public assemblies for *D. anananassae* [DanaRS2.1 (Zhang et al.unpublished, NCBI)], *D. elegans* [Dele_2.0 (Richards et al. unpublished, NCBI)], *D. erecta* [DereRS2 (Zhang et al.unpublished, NCBI)], *D. pseudoobscura* [UCI_Dpse_MV25 (Liao et al. unpublished, NCBI)], *D. simulans* [Prin_Dsim_3.0 (Pinharanda et al. unpublished, NCBI)], *D. suzukii* [LBDM_Dsuz_2.1.pri (Paris et al. unpublished, NCBI)], and *D. yakuba* [Prin_Dyak_Tai18E2_2.0 (Reilly et al. unpublished, NCBI)]. For instance, for a gene family restricted to the *melanogaster* complex (*D. melanogaster*, *D. sechellia* and *D. simulans*), any species outside this complex (i.e., *D. ananassae*, *D. elegans*, *D. erecta*, *D. pseudoobscura*, *D. suzukii* and *D. yakuba*) was considered an outgroup. Thus, for each gene family having orphan candidates, Blastn searches were applied to search the syntenic genomic regions of the outgroup genomes for homologous sequences (bit score > 39 or identities > 60%). The found homologous syntenic sequences showing evidence of being transcribed (i.e., evidence from RNA-Seq alignment data) were searched—employing Blastp searches—for the absence of homologous open reading frames (bit score < 39 and positives < 60%).

## Supporting information

supplementary references

## Acknowledgments

We thank Nicolás Moreyra and Agustina Sztyrle for comments that helped in different stages of the present study, and Joanna Chiu (University of California, Davis) for providing access to the *D. suzukii* 2.0 genome. This work was supported by the Agencia Nacional de Promoción Científica y Técnica through grants awarded to JH and EH.

## Authors’ Contributions

JH conceived and designed the study, compiled and analyzed the data, and took the lead in writing the manuscript. FCA was involved in planning the work and analysis design; she also estimated rates of molecular evolution and gene gain/loss. SAB performed functional annotations and designed the figures. SR helped integrate genomic information and predict protein sequences. EH was involved in planning the work and supervised the project. All authors discussed the results and contributed to the final manuscript.

## Supplementary Material

Table S1. List of *D. melanogaster* seminal genes. As KSGs we included genes encoding proteins previously confirmed to be transferred by males into females during mating, those meeting stringent multiple criteria that indicate so according to Sepil et al. (2019), or those expressed in male reproductive tissues more than in any other tissue (according to modENCODE and FlyAtlas2) also encoding secretable proteins found in the mating plug [according to Avila et al. (2015) and Wigby et al. (2020)]. As candidates, we included our novel candidates as well as previously predicted seminal genes. We excluded genes expressed specifically in the testes (according to FlyAtlas2) that encode sperm proteins (Wigby et al. 2020), those candidates proposed only by Wigby et al. (2020) that show low expression in male reproductive tissues and higher expression in other male and female tissues (according to modENCODE and FlyAtlas2), and those proposed only by Ayroles et al. (2011) that do not encode secretable proteins (signalP). The evaluated conditions for the expression/secretion criterion and sources that previously identified the gene as seminal are shown for each gene (see supplementary references).

Table S2. SFPs of the *melanogaster* subgroup, the *virilis-repleta* radiation, tephritids, and mosquitoes. Sources and methods used to compile the list are summarized for each considered species (see supplementary references).

## Data Availability

Despite no new data were generated in support of this research, the compiled information and data underlying our analyses are available in the article, in its online supplementary material, and/or at the open-access databases duly mentioned in the text.

